# Mid-infrared optoacoustic microscopy with label-free chemical contrast in living cells and tissues

**DOI:** 10.1101/270082

**Authors:** Miguel A. Pleitez, Asrar Ali Khan, Josefine Reber, Andriy Chmyrov, Markus R. Seeger, Stephan Herzig, Marcel Scheideler, Vasilis Ntziachristos

**Affiliations:** Institute of Biological and Medical Imaging, Helmholtz Zentrum München, Neuherberg, Germany; Chair of Biological Imaging, Technische Universität München, München, Germany; Institute for Diabetes and Cancer, Helmholtz Zentrum München, Neuherberg, Germany; Joint Heidelberg-IDC Translational Diabetes Program, Heidelberg University Hospital, Heidelberg, Germany; Molecular Metabolic Control, Medical Faculty, Technische Universität, München, Germany; Germany Center for Diabetes Research (DZD), Neuherberg, Germany

## Abstract

We developed mid-infrared optoacoustic microscopy (MiROM), a bond-selective imaging modality that overcomes water/tissue opacity and depth limitations of mid-infrared sensing allowing uncompromised live-cell/thick-tissue mid-infrared microscopy with up to three orders of magnitude higher sensitivity than other vibrational imaging modalities; such as Raman. We showcase the functional label-free biomolecular imaging capabilities of MiROM by monitoring the spatiotemporal dynamics of lipids and proteins during lipolysis in living adipocytes. Since MiROM, contrary to Raman modalities, is not only able to detect lipids and proteins, but also important metabolites such as glucose without the need of labels, here we discuss how MiROM yields novel functional label-free abilities for a broader range of analytical studies in living cells and tissues.

---

Label-free dynamic imaging of biomolecules (*i.e.* lipids, proteins, carbohydrates, and nucleic acids) in living cells remains challenging for optical microscopy. Visible (**VIS**) light microscopy cannot detect intrinsic biomolecular contrast directly, so it relies mainly on contrast provided by endogenous or exogenous chromophores. Ultraviolet (**UV**) microscopy detects nucleic acids, proteins, and lipids at high spatial resolutions, but at the expense of cell damage due to elevated photocytotoxicity. Additionally, cell auto-fluorescence has proven to be too weak for reliable live-cell imaging and, therefore, fluorescence microscopy depends mainly on the use of exogenous labels. Nonlinear microscopy modalities such as second-and third-harmonic generation can be used to image protein and lipids in cells and tissues, however the contrast provided here depends on the spatial organization and optical heterogeneity of the imaged structures but not on their chemical composition. The limitations of optical microscopy for label-free biomolecular contrast at wavelengths below 700 nm are fundamentally based on the chemically nonspecific nature of the electronic transitions and the threats of photodamage.

Chemically specific vibrational imaging modalities such as Coherent Raman Scattering (**CRS**) and mid-infrared (**mid-IR**) microscopy have enormously extended the range of possibilities for endogenous biomolecular imaging^1–4^. Stimulated Raman Scattering (**SRS**) microscopy, for instance, has been demonstrated in cells and tissues at high spatial resolution^5–7^. However, Raman-based modalities carry high risk of photodamage and offer detection limits above 1 mM^1^, which is inadequate for measuring micro-and nanomolar concentrations of target molecules in live cells. Direct vibrational excitation in the mid-IR range, on the other hand, provides cross-sections up to eight orders of magnitude larger than Raman methods, leading to higher sensitivity that can detect low-micromolar concentrations^8^. Microspectroscopy in the mid-IR range, for instance, can even allow rapid cancer diagnosis from histological samples^2,9–11^. Nevertheless, the application of mid-IR microscopy on live-cell imaging has been highly restricted mainly because water strongly absorbs light in this spectral range. Water is vital for cell survival and function, but at the same time its presence renders cell cultures opaque to mid-IR radiation. To reduce interfering water, samples can be placed within cuvettes approximately 10 µm thick and irradiated with high-powered mid-IR quantum cascade lasers (**QCL**s)^12,13^ or mid-IR synchrotron sources^14,15^. However, such confinement perturbs normal cellular behaviour and proliferation and, therefore, these alternatives have not been broadly implemented for live-cell imaging.

Here we introduce mid-IR optoacoustic microscopy (**MiROM**), a bond-selective imaging modality that detects specific vibrational transitions of biomolecules based on radiation-less deexcitation mechanism for highly efficient optoacoustic generation. We hypothesized that MiROM could bring high signal to ratio (SNR) label-free detection of biomolecules, addressing limitations of current label-free microscopy methods. Ultrasound-based detection of optical contrast circumvents the limitations of mid-IR light attenuation in water. Therefore, MiROM provides the chemical specificity of mid-IR sensing without opacity limitations due to water and without confining the sample to a thin cuvette. To capitalize on these advantages, MiROM was herein implemented in transmission-mode. This configuration allowed diffraction limited focused optical excitation with coaxially focused ultrasound detection, with excitation and detection sharing the same focal plane (**figure 1 a,b**). An absorption-contrastmicrograph is yielded by raster-scanning the sample within the focal plane while simultaneously acquiring the optoacoustic signal, the latter propagating without strong attenuation through long paths in water. Biomolecular specificity is achieved by tuning the emission of a broadly-tunable pulsed QCL to wavelengths corresponding to specific vibrational transitions.

**Figure 1:**
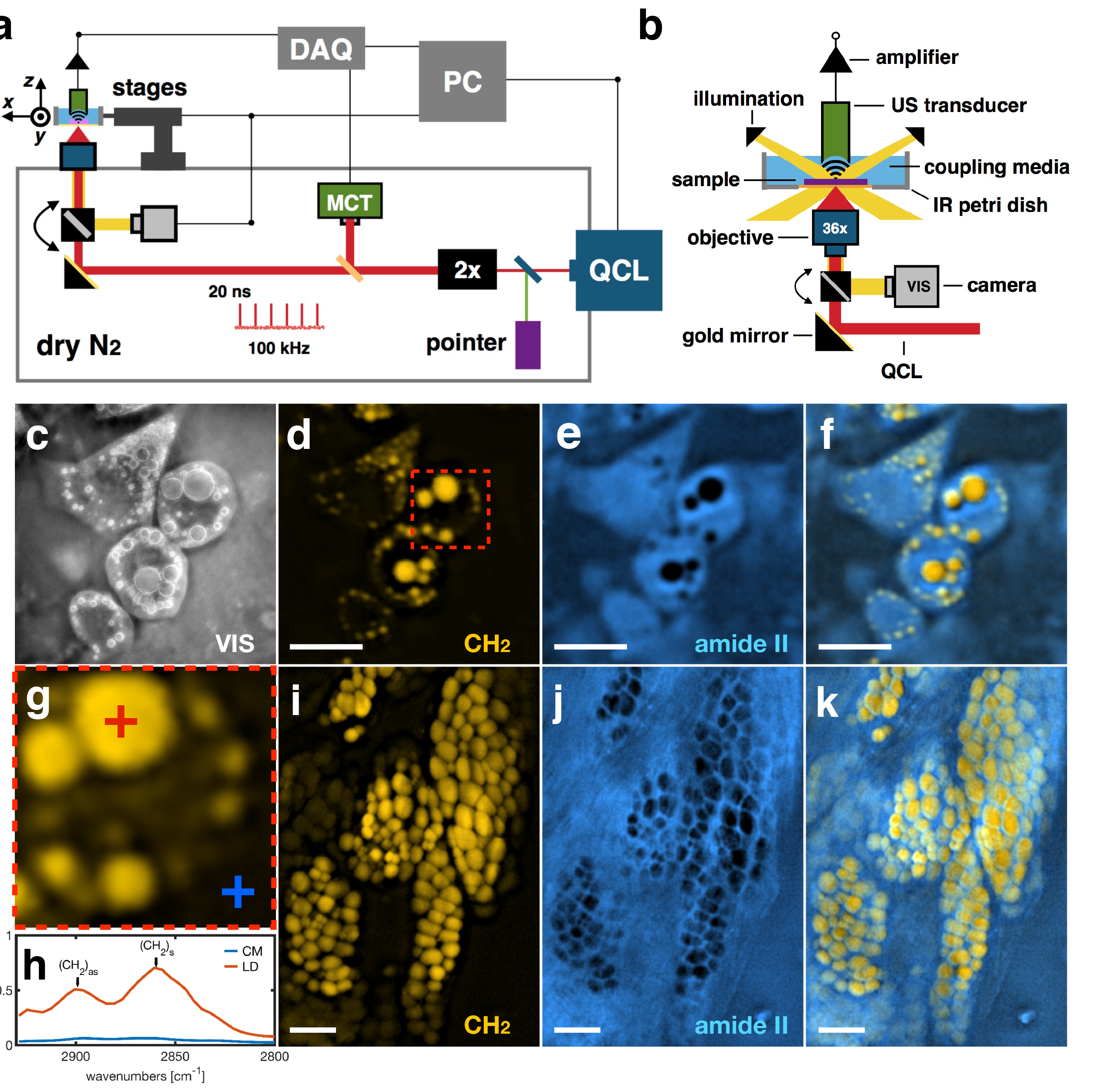
Mid-IR optoacoustic microscopy (MiROM). **(a)** General diagram of the imaging system. A tunable pulsed Quantum Cascade Laser (**QCL**) provides the excitation for optoacoustic imaging. An absorption intensity map is obtained by displacing the sample along the focal plane while simultaneously acquiring the optoacoustic signal. **(b)** excitation-sample-sensor configuration for MiROM. The focused ultrasound transducer and the reflective objective are coaxially aligned to share the same focal plane where the sample is placed (**see methods**). **(c)** VIS light micrograph of differentiated3T3-L1 cells, for reference. **(d-f)** MiROM micrographs of the cells in **c** with endogenous contrast for lipids **(d)** at 2857 cm^−1^ (CH_2_ vibration) and for proteins **(e)** at 1550 cm^−1^ (amide II). **(f)** merged lipid and protein maps. The FOV of the micrographs **d-f** is 175 µm × 175 µm, the scale bar is 50 µm. **(g)** zoom of a single adipocyte; dashed red square in **d**. Two spots for spectral analysis and fine tuning of the imaging wavelengthhave been marked; on a lipid droplet (**LD**) (red cross) and on the cell media (**CM**) (blue cross). **(h)** optoacoustic spectra in the CH vibrational region for the selected spots in **g**. **(i-k)** MiROM micrographs of freshly excised unstained pancreatic mouse tissue (C57BL/6) of 4 mm thickness, FOV of 550 µm × 800 µm, the scale bar is 100 µm. **(i)** lipid map at 2850 cm^−1^. **(j)** protein map at 1550 cm^−1^. **(k)** merged lipid and protein maps. Here, clusters of pancreatic acinar glands embedded in protein are observed.

We demonstrated the bond-selective live-cell imaging capabilities of MiROM by mapping the lipid and protein content of differentiated 3T3-L1 adipocytes. Endogenous contrast was detected by excitation of the symmetric CH_2_ vibration of lipids at 2857 cm^−1^ and excitation of the amide II band of proteins at 1550 cm^−1^, mainly from NH bending and CN stretching. The spatial resolution of the system (about 5.3 µm at 2850 cm^−1^, **supplementary figure 1**) allowed the identification of single adipocytes as well as individual lipid droplets (**LD**s) (**figure 1 c-g**). In the micrograph obtained at the CH_2_ vibration (the lipid map), LDs offered strong positive contrast due to their high content of triglycerides compared to the cell body and background cell medium (**CM**); here contrast-to-background ratios (**CBR**s) up to 14:1 were observed, while the signal-to-noise ratios (**SNR**s) were as high as 223:1 (see **methods** for details). In the micrographs obtained at the amide II band (the protein map), positive contrast came mainly from the overall protein content of the cells as well as from a weak contribution from water. Here, CBRs up to 1.7:1 and SNRs around 80:1 were observed (**supplementary figure 2**). In the protein map, LDs offered negative contrast because they are hydrophobic and contain much less protein than the cytoplasm. The absolute contrast of LDs in the protein map was nevertheless high enough to be accurately detected (SNR up to 46:1), perhaps reflecting the content of lipid droplet-associated proteins (**supplementary figure 2**). Additionally to lipid and protein maps, carbohydrate contrast was obtained on 3T3-L1 adipocytes by excitation at 1081 cm^−1^ (C - O deformation) assigned to carbohydrates. Localized contrast at this wavelength was found on the cell body around the growing lipid droplets (**supplementary figure 3**). One possible explanation for the presence of carbohydrates on the cell body might be the capture and accumulation of glucose involved in the synthesis of triglycerides to be packed into the lipid-droplets during lipogenesis.

Since established opaque-sample mid-IR sensing techniques, such as reflectometry and Attenuated Total Reflectance (**ATR**), cannot extract information from depths beyond 2 µm^2,16^, we investigated whether MiROM can be applied to mm-thick tissues. Here, the penetration depth is only limited by the absorption coefficient, which in tissues can be up to 100 µm depending on water content. This was done by mapping the lipid content (at 2850 cm^−1^) and protein content (at 1550 cm^−1^) of a 4-mm-thick slice of freshly excised pancreatic tissue from a C57BL/6 mouse (**figure 1 i-k**). No histology preparation was performed (see **methods** for details). In this case, the pancreatic acinar glands are seen in positive contrast in the lipid map (CBR up to 44:1) while the protein map shows, in negative contrast, the compartments where the acinar glands are embedded (CBR up to 6:1) (**supplementary figure 2**). The CBRs for lipid and protein maps were at least 3.5 times higher with thick tissues than with monolayer cell cultures in aqueous medium (**figure 1 d-f**).

Given our ability to obtain high SNRs even with averaging times of only 1 ms/pixel, we used MiROM to monitor the dynamics of lipids and proteins during isoproterenol-induced lipolysis in white adipocytes (differentiated 3T3-L1) and brown adipocytes (differentiated PreBAT) over a period of 4 hours. Lipid maps (at 2857 cm^−1^) and protein maps (at 1550 cm^−1^) were taken every 5 minutes before and after addition of isoproterenol to the medium (see **methods** for details). In both adipocyte types, overall lipid content increased at a low rate before lipolysis induction, reflecting ongoing lipogenesis. After induction, lipid contrast clearly decreased continuously and nearly linearly with time (**figure 2 a-f**). Different adipocytes exhibited different lipolysis rates (**figure 2e,f**), with some adipocytes unaffected by isoproterenol. Continuous LD remodeling, when larger LDs absorbed smaller ones, was observed in some adipocytes (**figure 2 a, ROI 1**). Other adipocytes became notably dimmer with time after lipolysis induction (**figure 2 a**, **ROI 2**). The same heterogeneous response to isoproterenol was observed in brown adipocytes (**figure 2 b,f**). Lipolysis was faster and more extensive in brown adipocytes than in white adipocytes: by 2 hours, lipid contrast had changed up to 30% in brown adipocytes, compared to 18% in white adipocytes. (see also **supplementary videos V1 and V2**).

**Figure 2:**
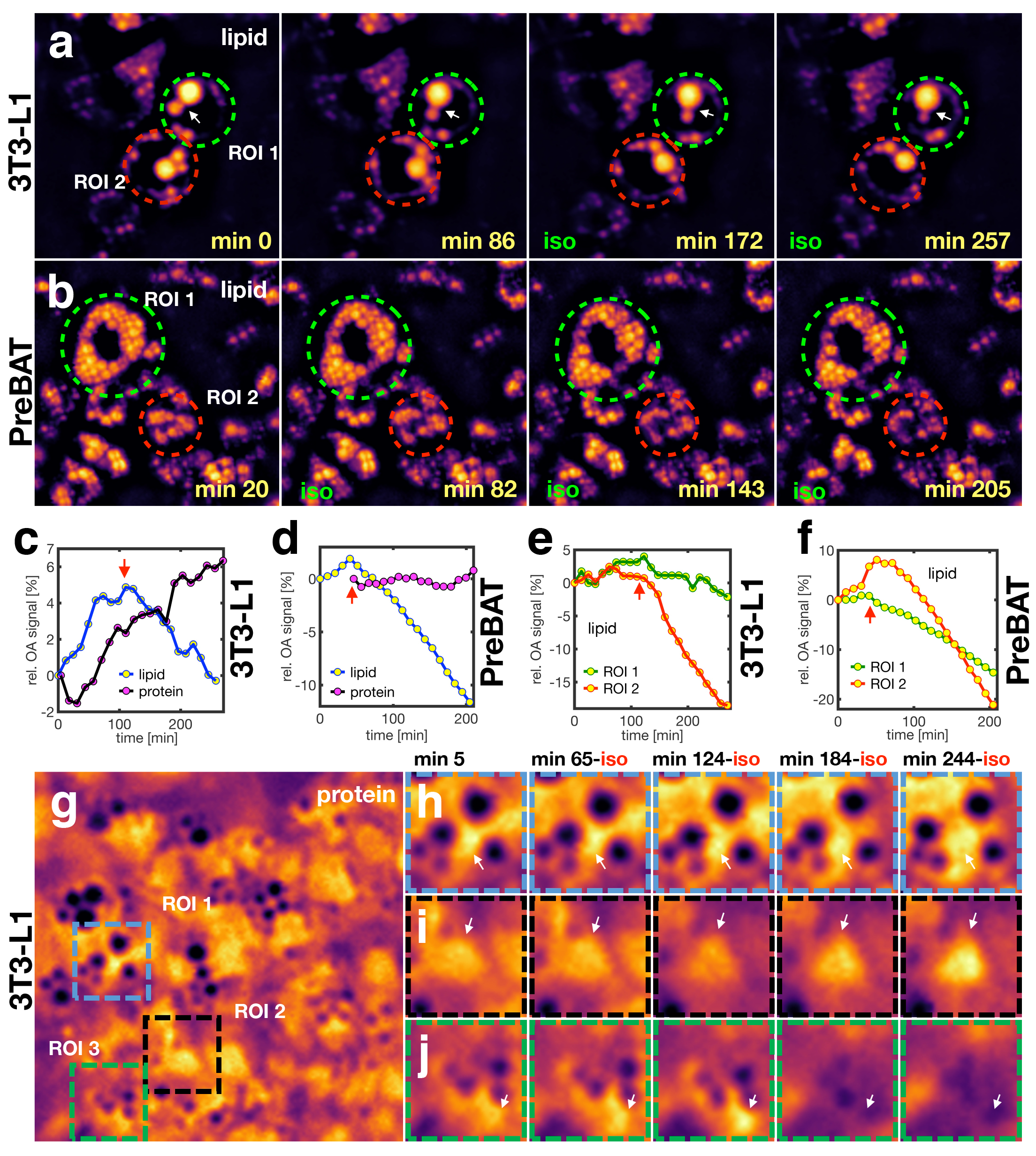
Functional live-cell mid-IR optoacoustic microscopy. Monitoring of induced lipolysis in **(a)** white (differentiated 3T3-L1) and **(b)** brown 15(differentiated PreBAT) adipocytes;only lipid maps at 2857 cm^−1^ are shown (see also **supplementary figure 4** for protein/merged maps). Two regions-of-interest (**ROI**) enclosing individual adipocytes are marked on each sequence, green dashed circle for ROI 1 and red dashed circle for ROI 2. The white arrow in **a** follows the process of lipid droplet remodeling observed in the single white adipocyte enclosed in ROI 1. Time and presence of ISO is indicated at the bottom corners of each frame. Overall relative lipid and protein contrast change for **(c)** white and **(d)** brown adipocytes. Relative lipid contrast change for ROIs 1 and 2 in **(e)** white and **(f)** brown adipocytes. Thered arrow in **c-f** indicates when ISO was added. **(g)** protein map (at 1550 cm^−1^) of white adipocytes (a different batch tothe one in **a**). Here, three ROIs have been marked: blue dashed square for ROI 1, black dashed square for ROI 2, and green dashed squarefor ROI 3. Each ROI is centered on a single adipocyte (see also **supplementary images 5** for lipid/merged maps). **(h-j)** Monitoring of proteins during lipolysis for the selected ROIs in **c**. The white arrow follows regions changing protein contrast during lipolysis. Common time and presence of ISO for each ROI, is indicated above sequence **h**. The FOV of **b** and **g** is250 µm × 250 µm; the batch of cells and FOV in **a** is the same as in **figure 1 d-f**.

Even more dynamic and heterogeneous behavior was observed when monitoring the protein content of individual cells during lipolysis (**figure 2 g-j**). Some protein-containing areas expanded over time (**figure 2 h**), others showed decreased contrast followed by partial recovery (**figure 2 i**), and some areas increased in protein contrast, then showed reduced contrast with no recovery (**figure 2 j**; see also **supplementary videos V3**). We hypothesize that these heterogeneous changes of protein contrast in response to lipolysis induction may reflect a combination of adipokine secretion, protein degradation, and/or protein translation. Additionally, since the amide II band is conformation-dependent^17,18^, it is also possible that the changes in protein contrast reflect ongoing conformational changes in cell proteins.

MiROM offers unprecedented high contrast, image quality, and sensitivity for mid-IR microscopy of living cells as well as for thick unprocessed tissues, avoiding at the same time the problems of scattering and speckles common when using highly coherent mid-IR sources; unlike, for instance, mid-IR photothermal microscopy^19–21^. It not only complements other vibrational imaging platforms, such as Raman-based modalities, where the amide II band is practically silent^18^, but also offers extended molecular contrast since, contrary to Raman, is also able to detect carbohydrates in living cells. Furthermore, MiROM can potentially offer greater sensitivity than Raman methods because it is based on direct vibrational excitation. For instance, here, we achieved spatiotemporal monitoring of intrinsic lipid/protein contrast changes of <1% during lipolysis in living adipocytes. To our knowledge, this has not been previously reported with Raman or other microscopy modalities based on intrinsic molecular contrast.

Finally, although here we applied MiROM for the assessment of lipids, proteins, and carbohydrates, the same can also be applied to detect, nucleic acids, and water in practically any other cell culture or tissue. In this way, MiROM may become an important tool in live-cell biology and analytical histology, while filling an important gap in vibrational imaging and considerably extending the contrast portfolio of optoacoustic microscopy previously limited only between the UV and the near infrared spectral region.

## Methods

### System description

A broadly tunable pulsed Quantum Cascade Laser (**QCL**) (MIRcat, Daylight Solutions, CA, USA) is used for optoacoustic generation and biomolecular specificity; the spectral range of the QCL is 3.4 - 11.0 µm (2941- 909 cm^−1^). The pulse duration of the QCL is set to 20 ns at a repetition rate of 100 kHz then focused into the sample by a 0.5 NA reflective objective (36x, Newport Corporation, CA, USA). The mid-IR absorption map of the sample is obtained by displacing the sample along the focal plane by motorized stages (Prior Scientific, Cambridge, UK) simultaneously detecting the optoacoustic signal by a 20 or a 25 MHz central frequency focused ultrasound transducer (Imasonic/Sonaxis, Voray sur l’Ognon/Besancon, France). The focused ultrasound transducer and the reflective objective are coaxially aligned to share the same focal plane where the sample is placed on a custom-made mid-IR transparent stainless-steel petri-dish using a ZnSe window (Edmund Optics, Mainz, DE) as bottom substrate. For the live-cell studies, the cell media served as acoustic coupling between the US transducer and the cells. Deionized water was used as coupling media otherwise.

To remove interference from atmospheric CO_2_ and water vapor, the mid-IR beam-path is purged with a constant flow of dry N_2_. A Mercury-Cadmium-Tellurium (**MCT**) detector (Daylight Solutions, CA, USA) is used for optical reference and a VIS laser pointer, co-aligned with the QCL beam, serves as aiming beam for easy optical adjustment (see **figure 1 a**). For validation, co-registration, and easy ROI selection, oblique VIS illumination (Edmund Optics, Mainz, DE) is used to obtain standard light micrographs with a general purpose monochromatic camera (Edmund Optics, Mainz, DE) (see **figure 1 b**). Prior to measurements on cells and tissues, the contrast and resolution of the system was tested imaging synthetic samples; polyamide sutures and polyethylene microspheres embedded in agar, (**Supplementary figure 1**).

### Signal recovery and Signal-to-Noise Ratio

The raw optoacoustic signals were amplified by 63 dB (MITEQ, NY, USA) filtered with at 50 MHz low pass filter (Mini-Circuits, NY, USA) and then recorded at a sampling rate of 250 MS/s on a 12 bit DAQ card (Gage Applied, Lockport, USA) The intensity of each pixel composing the micrographs shown in this work is the peak-to-peak amplitude value resulting from the average of 100 optoacoustic transients; corresponding to a pixel measurement time of 1 ms at the pulse repetition rate of 100 kHz.

The signal-to-noise ratio is defined here as the ratio between the peak-to-peak amplitude value of the optoacoustic signal vs. the peak-to-peak amplitude value of the noise level before the arrival of the optoacoustic signal. For instance, the maximum SNR of the lipid map for the white adipocytes discussed above is 223, corresponding to a relative error of 0.45 %. In terms of absolute values this corresponds to an optoacoustic peak-to-peak amplitude of 702.2 mV and a peak-to-peak amplitude of the noise of 3.1 mV. In the protein map the maximum SNR is close to 80, or 1.3 % relative error. This is calculated from an absolute optoacoustic peak-to-peak amplitude of 247 mV and a noise level of around 3 mV.

### Image processing

In order to enhance visibility and compensate for spatial resolution of mid-IR microscopy in the range of subcellular compartments of interest (~ 5 µm), the images were bicubic interpolated to a pixel size of 250 nm and deconvolved with the experimental determined point-spread function (**see supplementary fig. 1**) by a 3 or 5 step iterative Wiener deconvolution. Furthermore, the images were post processed by a 2-pixel Gaussian filtering, outlier removal if necessary, a contrast enhancement to a 0.3% saturation, and a histogram normalization. For resolution analysis and SNR determination, the images were kept unprocessed.

### Preparation and measurement of white and brown adipocytes

3T3-L1 mouse white preadipocyte cells were plated on the custom plates and cultured in growth media containing DMEM containing 1 g/l glucose (Life Technologies, Paisley, Scotland, 10% FBS (Merck, Darmstadt, DE) and 1% Pen/Strep (Life Technologies, Bleiswijk, Netherlands)) till confluency. The process of differentiation lasted for 6 days. On day 0 and day 2, the growth media with 1 μg/ml insulin (Sigma, Steinheim, DE), 0.25 uM dexamethaxone (Sigma, Steinheim, DE), 0.5 mM 3-isobutyl-1-methylxanthine (Sigma, Steinheim, DE) and 1/1000 volume ABP (50 mg/ml L-ascorbate, 1 mM biotin, 17 mM pantothenate; Sigma, Steinheim, DE) was added to the cells. On day 4 of differentiation, growth media with only insulin and ABP was used while on day 6 only growth media was added.

The PreBAT cell line was created and provided by Hoppmann, Perwitz et al. (2010) by immortalizing pre-adipocytes from the intrascapular BAT of newborn mice using the SV40 Large T antigen. The differentiation process also lasted for 6 days starting with induction on day 0 with DMEM growth media (4.5 g/l glucose; Life Technologies, Paisley, Scotland) containing 20 % FBS (Merck, Darmstadt, DE) and 1 % Pen/Strep (Life Technologies, Bleiswijk, Netherlands) with the addition of 20 nM insulin (Sigma, Steinheim, DE), 1 µM triiodothyronine (T_3_) (Sigma, Steinheim, DE), 0.125 mM indomethacin (Sigma, Steinheim, DE), 2 µg/ml dexamethasone (Sigma, Steinheim, DE) and 0.5 mM 3-isobutyl-1-methylxanthine (Sigma, Steinheim, DE). On day 02 and day 04, growth media containing only insulin and T_3_ was added and on day 06 the cells were given only growth media. At the end of differentiation, both cell lines showed abundant amounts of lipid droplets.

### Preparation and measurement of mouse tissues

Male C57BL/6J mice (8-10 weeks old; Charles River Laboratories Inc, Charleston, USA) were kept at 24±1°C and fed with standard rodent diet (Altromin 1314, Altromin Spezialfutter GmbH & Co, Germany) with free access to water, with constant humidity and on a 12-h light-dark cycle. After the mice were sacrificed the organs were harvested and directly placed on the sample holder where they were covered with low temperature melting agar (2%) and de-ionized (DI) water as coupling media.

## Acknowledgements

The research leading to these results has received funding from the Deutsche Forschungsgemeinschaft (DFG), Germany [Gottfried Wilhelm Leibniz Prize 2013; NT 3/10-1], as well as from the European Research Council (ERC) under the European Union’s Horizon 2020 research and innovation programme under grant agreement No 694968 (PREMSOT).

## Author Contributions

**M.A.P.** created the imaging concept, designed, built, and characterized the imaging system. **M.A.P.**, **A.A.**, and **J.R.** designed and performed the experiments on adipocytes. **M.A.P.** and **J.R.** designed and performed the experiments on excised tissues. **A.C.** synchronized and automated the imaging system. **M.R.S.** performed the image processing and prepared the art work. **M.A.P.** processed the results, prepared the images, and wrote the manuscript. **M.A.P.**, **A.A.**, **J.R.**, and **M.S.** analyzed the results on lipolysis. **M.S.** and **S.H.** supervised the study on lipolysis. **V.N.** supervised the whole study. All authors edited the Manuscript.

## Bibliography

1. Cheng, J.-X & Xie, X. S. Vibrational spectroscopic imaging of living systems: An emerging platform for biology and medicine. Science (80-.). 350, aaa8870–aaa8870 (2015).

2. Baker, M. J. et al. Using Fourier transform IR spectroscopy to analyze biological materials. Nat. Protoc. 9, 1771–91 (2014).

3. Bhargava, R. Infrared Spectroscopic Imaging: The Next Generation. Appl. Spectrosc. 66, 1091–1120 (2012).

4. Diem, M.et al. Molecular pathology via IR and Raman spectral imaging. J. Biophotonics 6, 855–886 (2013).

5. Prince, R. C.Frontiera, R. R. & Potma, E. O. Stimulated Raman Scattering: From Bulk to Nano. Chem. Rev. 117, 5070–5094 (2017).

6. Li, J. & Cheng, J.-X. Direct Visualization of De novo Lipogenesis in Single Living Cells. Sci. Rep. 4, 6807 (2014).

7. Wei, L. et al. Super-multiplex vibrational imaging. Nature 544, 465–470 (2017).

8. Schwaighofer, A., Alcaráz, M. R., Araman, C. & Goicoechea, H. External cavity-quantum cascade laser infrared spectroscopy for secondary structure analysis of proteins at low concentrations. Nat. Publ. Gr. 1–10(2016).doi:10.1038/srep33556

9. Byrne, H. J. et al. Spectropathology for the next generation: Quo vadis? Analyst 140, 2066–2073 (2015).

10. Walsh, M. J.,Reddy, R. K. & Bhargava, R. Label-free biomedical imaging with mid-IR spectroscopy. IEEE J. Sel. Top. Quantum Electron. 18, 1502–1513 (2012).

11. Bird, B. et al. Infrared spectral histopathology (SHP): a novel diagnostic tool for the accurate classification of lung cancer. Lab. Investig. 92, 1358–1373 (2012).

12. Haase, K., Kröger-Lui, N., Schönhals, A. &Petrich, W. Real-time mid-infrared imaging of living microorganisms. J. Biophotonics 9, 61–6 (2016).

13. Schwaighofer, A. Brandstetter, M., & Lendl, B. Chem Soc Rev Quantum cascade lasers (QCLs) in biomedical spectroscopy His work focuses on applying. Chem. Soc. Rev. 46, 5903–5924 (2017).

14. Holman, H.-Y. N., Bechtel, H. A., Hao, Z. & Martin, M. C. Synchrotron IR Spectromicroscopy: Chemistry of Living Cells. Anal. Chem. 82. 8757–8765 (2010).

15. Martin, M. C. et al. 3D spectral imaging with synchrotron Fourier transform infrared spectro-microtomography. Nat. Methods 10, 861–4 (2013).

16. Shaw, R. A. & Mantsch, H. H. Infrared Spectroscopy in Clinical and Diagnostic Analysis. Encycl. Anal. Chem. 1–20 (2006). doi:10.1002/9780470027318.a0106

17. Oberg, K. A., Ruysschaert, J.-M. & Goormaghtigh, E. The optimization of protein secondary structure determination with infrared and circular dichroism spectra. Eur. J. Biochem. 271, 2937–2948 (2004).

18. Barth, A. Infrared spectroscopy of proteins. Biochim. Biophys. Acta 1767, 1073–101 (2007).

19. Lee, E. S. & Lee, J. Y. High resolution cellular imaging with nonlinear optical infrared microscopy. 19, 1378–1354 (2011).

20 Zhang, D. et al. Depth-resolved mid-infrared photothermal imaging of living cells and organisms with submicrometer spatial resolution. Sci. Adv. 2, (2016).

21 Bai, Y., Zhang, D., Li, C., & Cheng, J.-X. Bond-Selective Imaging of Cells by Mid-Infrared Photothermal Microscopy in High Wavenumber Region. J. Phys. Chem. B acs.jpcb.7b09570 (2017). doi:10.1021/acs.jpcb.7b09570

